# *Hox* genes pattern the primary body axis of an anthozoan cnidarian prior to gastrulation

**DOI:** 10.1101/219758

**Authors:** Timothy Q. DuBuc, Thomas B. Stephenson, Amber Q. Rock, Mark Q. Martindale

## Abstract

*Hox* gene transcription factors are important regulators of positional identity along the anterior-posterior axis in bilaterian animals. Cnidarians (e.g. sea anemones, corals and hydroids) are the sister group to the Bilateria and possess genes related to both anterior and central/posterior class *Hox* genes. In the absence of a conserved set of *Hox* genes among other early branching animal clades, cnidarians provide the best opportunity to learn about the emergence of this gene family. We report a previously unrecognized domain of *Hox* expression in the starlet sea anemone, *Nematostella vectensis*, beginning at early blastula stages. Functional perturbation reveals that two *Hox* genes not only regulate their respective expression domains, but interact with one another to pattern the entire oral-aboral axis mediated by Wnt signaling. This suggests an ancient link between *Hox*/*Wnt* patterning of the oral-aboral axis and suggest that these domains are likely established during blastula formation in anthozoan cnidarians.

---

*Hox* genes are a specific family of homeobox-containing transcription factors that have been studied extensively in several clades of bilaterally symmetrical animals (“bilaterians”, Fig. 1a). Originally discovered in *Drosophila*^1^, *Hox* genes play an important role in establishing segment identity along the anterior-posterior (A-P) axis during development, and are conserved in all bilaterian lineages including vertebrates^2^. *Hox* genes are often clustered together along contiguous stretches of an animal’s genome^3^, and are classified into three paralogy groups: anterior (*Hox*1-3), central (*Hox*4-8) and posterior (*Hox*9-13) (Fig. 1b). Early comparisons of vertebrate and insect *Hox* sequences revealed that homologous *Hox* genes occupy similar locations in the 3’-5’ topology of the cluster^4^ and gene expression analysis further revealed that a gene’s position in the cluster was correlated to its expression profile along the anterior-posterior axis. Although there are derivations in the organization and distribution of *Hox* genes among bilaterians, *Hox* genes are consistently expressed in anterior-to-posterior territories as reflected by their phylogenic ancestry^5–15^. Furthermore, duplications and fragmentation of *Hox* clusters likely had a strong impact on the radiation of vertebrates,^9,16–18^ although the origin and function of this gene family within the first invertebrates is less clear^19–22^.

**Figure 1:**
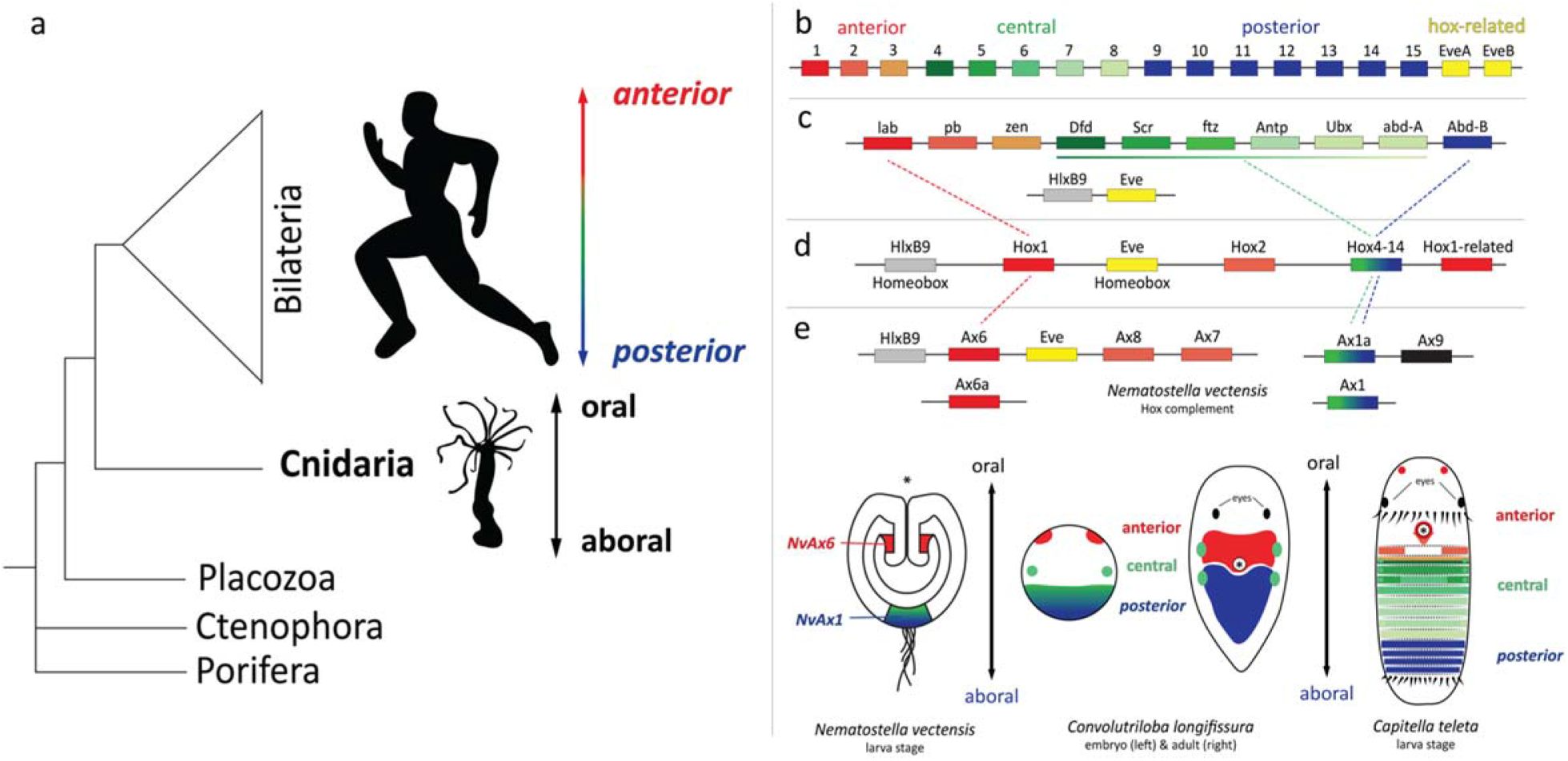
Anterior-posterior patterning and the emergence of a *Hox* cluster. **(a)** Bilaterians are classically defined by an anterior-posterior axis perpendicular to the dorsal ventral axis. Cnidarians are the sister taxa to bilaterians and are the only basal lineage to have a diverse cluster of *Hox* genes. (**b**) The common ancestor of the deuterostome lineage likely had a *Hox* cluster consisting of 14-15 *Hox* genes, closely associated with the homeobox gene *Eve^18^*. (**c)** Evidence from the protostome, *Tribolium castaneum*, suggests that the protostome ancestor also had an intact *Hox* cluster consisting 10 linked *Hox* genes^24,77,78^. **(d)** The cnidarian ancestor had both anterior (*Hox1* and *Hox2*) and central/posterior (*Hox*9-13) class *Hox* genes^30^. **(e)** The *Hox* complement of the anthozoan cnidarian, *Nematostella vectensis*, has phylogenetically anterior (*NvAx6, NvAx6a, NvAx7* and *NvAx8*) and central/posterior (*NvAx1* and *NvAx1a) Hox* genes^20,21^. Depiction of *Hox* expression along the oral-aboral axis in diverse invertebrates. Regions of anterior, central and posterior *Hox* expression are designated with shades of red, green and blue respectively. The anterior (*NvAx6*) and central/posterior (*NvAx1) Hox* genes of *Nematostella* are expressed along the oral-aboral axis during larval development. (* = site of mouth formation).

Phylogenetic reconstruction of the bilaterian *Hox* gene compliment indicate that the common ancestor of most extant animals had a *Hox* cluster containing anterior, central, and posterior *Hox* genes^18,23,24^ (Fig. 1b-c). Although clustering is not an essential component of *Hox* functionality, it is widely associated with diverse animal clades from cnidarians and throughout Bilateria. Cnidarians (e.g. corals, anemones, and hydroids) are the sister group to the Bilateria^25,26^ (Fig. 1a) and possess bona fide *Hox* genes related to anterior and central/posterior *Hox* genes^20,21,27–31^. The cnidarian common ancestor likely had a *Hox* cluster consisting of both anterior and a representative of an ancestor to a clade sister to both central and posterior genes^30^ (Fig. 1d, Supplemental Fig. 1b). Definitive central class genes have yet to be found in a cnidarian, first appear in the Xenoacoelomorph bilaterian lineage^32^ and expanded in protostome/deuterostome lineages (Fig. 1e). Currently, it is uncertain whether central class *Hox* genes arose in the bilaterian lineage from an ancestral central/posterior *Hox* gene, or were lost in the cnidarian lineage. Bona fide *Hox* genes have yet to be found in the genomes of earlier branching animal clades although some evidence suggests a loss of this gene family^22,33–37^. Efforts to functionally characterize the cnidarian *Hox* complement are crucial for determining the pre-bilaterian role of these important developmental regulators.

Functional studies in the fly^1^ and mouse^38^ first showed that *Hox* genes are causally involved in establishing adult body structures from the region in which they are expressed, thus controlling regional identity along the A-P axis. *Hox* genes interact with one another in overlapping domains, with posterior *Hox* genes having functional “dominance” over more anterior genes, however flies also exhibit examples of anterior dominance^39–41^. mRNA expression studies of *Hox* genes in multiple cnidarian species suggest a discrete role in late larval patterning, yet the developmental function remains untested^19,21,28,29,42–45^. Here we begin to dissect the functional role and hierarchy of *Hox* genes in anthozoan cnidarian, *Nematostella vectensis*, and generate a presumptive molecular network for oral-aboral patterning that may have functioned in the cnidarian ancestor.

## Results

### *Hox* genes of *Nematostella* are expressed prior to germ layer segregation

The *Hox* cluster of the starlet sea anemone *Nematostella* consists of two neighboring non-Hox homeobox genes (*NvHlxB9 and NvEve*), along with putative orthologs to the anterior *Hox* genes *Hox1 (NvAx6*) and *Hox2 (NvAx7* and *NvAx8*)^20,21^. Four additional *Hox* genes exist in the genome, consisting of: two central/posterior genes (*NvAx1* and *NvAx1a*), a pseudogene (*NvAx9*), and an additional anterior-like gene of indeterminate orthology (*NvAx6a*) (Fig. 1e). The assemblage of *Hox* clusters from different cnidarian genomes indicate that anthozoan cnidarians had a relatively intact *Hox* cluster, which have been fragmented in different species (Supplemental Fig. 1). Corals arguably have the most complete cluster consisting of both anterior and central/posterior orthologs^30^ (Supplemental Fig. 1c). Representative anemone species (*Nematostella* and *Aiptasia*) appear to have diverged from a coral-like cluster with a duplicated *Hox2* ortholog (*Ax7* and Ax8) (Supplemental Fig. 1d-e). Additionally, the *Aiptasia* genes (Ax6 and Eve) are not on the same scaffold as the *Hox* cluster, yet the ortholog of *Ax6a* is found on the later part of the cluster^31^ (Supplemental Fig. 1e). Two orthologs of *Ax6a* were found linked in a coral^30^, but were not found linked to the other *Hox* genes (Supplemental Fig.1). In *Nematostella*, the pseudogene *NvAx9* is linked to *NvAx1a* and *Ax6a* and *Ax1a* are linked in *Aiptasia*. Therefore, we suspect that *NvAx9* is a derived Ax6a-related gene.

An ortholog of *NvAx1* is present in all cnidarian genomes, yet there is no evidence of it being linked to a cnidarian cluster, suggesting it diverged long ago. Genomes that exhibit a high rate of cluster fragmentation are correlated with an expansion of central/posterior genes and less conservation among anterior-like genes^20,21^ (Supplemental Fig. 1f). These findings are based on the idea that the ancestral condition was a state of *Hox* clustering, which is generally assumed to be the representative condition among bilaterian clades. A lack of functional data only allows us to speculate on the impact of cluster fragmentation within Cnidaria, and a more thorough assessment of the *Hox* complement across the phylum may help resolve the impact of the cnidarian *Hox* diversification.

Previous *in situ* hybridization studies conducted on larval and juvenile stages of *Nematostella* revealed that the majority of *NvHox* genes are expressed in a staggered domain along the primary (oral-aboral) axis^19, 20^ with the anterior *Hox* gene (*NvAx6*) expressed in a region corresponding to the pharyngeal nerve ring at the oral pole, and the central/posterior *Hox* gene (*NvAx1*) expressed in the larval apical tuft at the aboral tip of the planula (Fig. 1e). The rest of the *N. vectensis Hox* genes are expressed at planula stages asymmetrically along one side of the endoderm with an oral boundary intermediate between *NvAx6* and *NvAx1*^21,28^. These data along with other studies in medusazoan cnidarians that possess complex metagenic lifecycles have created uncertainty as to whether cnidarian *Hox* genes have any bilaterian-like attributes, and led to the suggestion that the axial patterning role of *Hox* genes arose after the cnidarian bilaterian split^19,29,44^.

We analyzed expression of *Hox* and neighboring non-Hox homeobox genes in *Nematostella* throughout early development (Fig. 2a), using a highly sensitive method (quantitative PCR) and verified expression by *in situ* hybridization. mRNA of both anterior and central/posterior *Hox* genes were detected early in development (Fig. 2b). Maternal mRNA of *NvAx1* and *NvAx6a* could be detected by qPCR in fertilized eggs (Fig. 2b, 0hpf), however transcriptional activation of the extended Hox gene cluster begins during early blastula formation (Fig. 2b, 12hpf) with expression of the *non-Hox* homeobox gene *NvHlxB9* and the adjacent anterior *Hox* gene *NvAx6*. Deployment of the *Hox* cluster continues during the blastula to gastrula transition, with the activation of *NvEve* (Fig. 2b, 16hpf) and two genes related to *Hox2 (NvAx7* and *NvAx8*) (Fig. 2b, 18hpf). Lastly, activation of the central/posterior *Hox* gene (*NvAx1a*), which is fragmented from the predicted ancestral cnidarian *Hox* gene cluster (Fig. 1d)^30^, begins near the onset of gastrulation (Fig. 2b, 24hpf).

**Figure 2:**
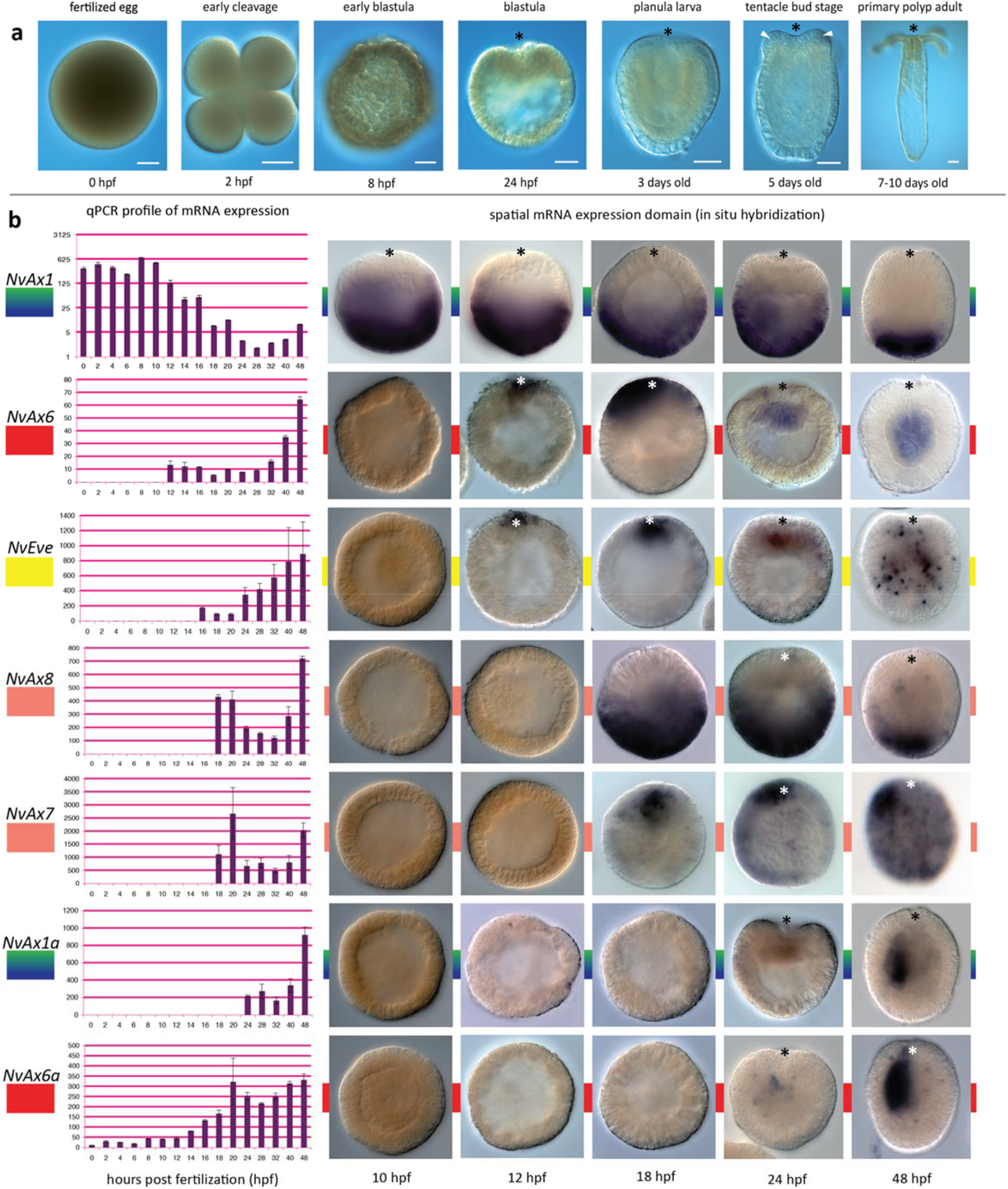
*Hox* genes of *Nematostella* exhibit hallmarks of bilaterian *Hox* genes during early development. **(a)** Time series of the embryonic, larval and adult development of the cnidarian, *Nematostella vectensis* (hours post fertilization=hpf). **(b)** Clustered *Hox* genes display temporal colinearity during early embryonic development. Expression begins with the 3’ located neighboring non-Hox homeobox gene *NvHlxB9* and the anterior gene *NvAx6* at twelve hpf. Subsequent activation of the other genes in the cluster maintains colinear expression relative to the ancestral cnidarian cluster. Two of the unlinked *Hox* genes *NvAx1* (central/posterior) and *NvAx6a* (anterior) appear maternally expressed during early development**. (c)** *In situ* hybridization confirms the temporal activation of the *Hox* cluster. Prior to gastrulation, mRNA expression of the central/posterior *Hox* gene (*NvAx1*) localizes to the aboral pole, while an anterior *Hox* gene (*NvAx6*) becomes transcriptionally activated along the oral pole. Remaining *Hox* and homeobox genes occupy oral (*NvEve, NvAv7, NvAx6a*) or aboral (*NvAx8*) domains during early development. Scale Bars are 50um.

Two zones of spatial activation were found, with genes being expressed in oral (*NvAx6, NvAx6a, NvEve, NvAx7*,) or aboral (*NvAx1, NvAx8*) domains before and during early gastrulation prior to any asymmetries along the directive axis (Fig. 2b). Notably, *NvAx6* and *NvAx1* were the first to be detected by *in situ* hybridization, occupying complimentary oral and aboral domains during early blastula formation (12hpf) (Fig. 2c, Supplemental Fig. 2). *NvHlxB9* and *NvEve* are expressed along the oral pole of the animal during early gastrulation,^46^ and in agreement with the qPCR analysis (Fig. 2b), the *NvHox* genes appear to be expressed along a similar temporal timeline. Respective oral and aboral expression of *NvAx6* and *NvAx1* continues through the blastula to gastrula transition. At this stage *NvAx6 is* initially expressed at the site of gastrulation (animal pole) in the presumptive endomesoderm before becoming restricted to the pharyngeal endoderm (Fig. 3a-e) associated with the pharyngeal nerve ring^21,28^. Conversely, *NvAx1* maintains a broad aboral expression domain throughout blastula and gastrula stages before becoming refined to the most aboral domain of the apical tuft region during early planula stages (Fig. 3f-j). The broad complimentary expression domains of the orally expressed anterior Hox gene *NvAx6* and aborally expressed *NvAx1* genes during blastula stages suggests they may play an important role in oral-aboral patterning during early development.

**Figure 3.**
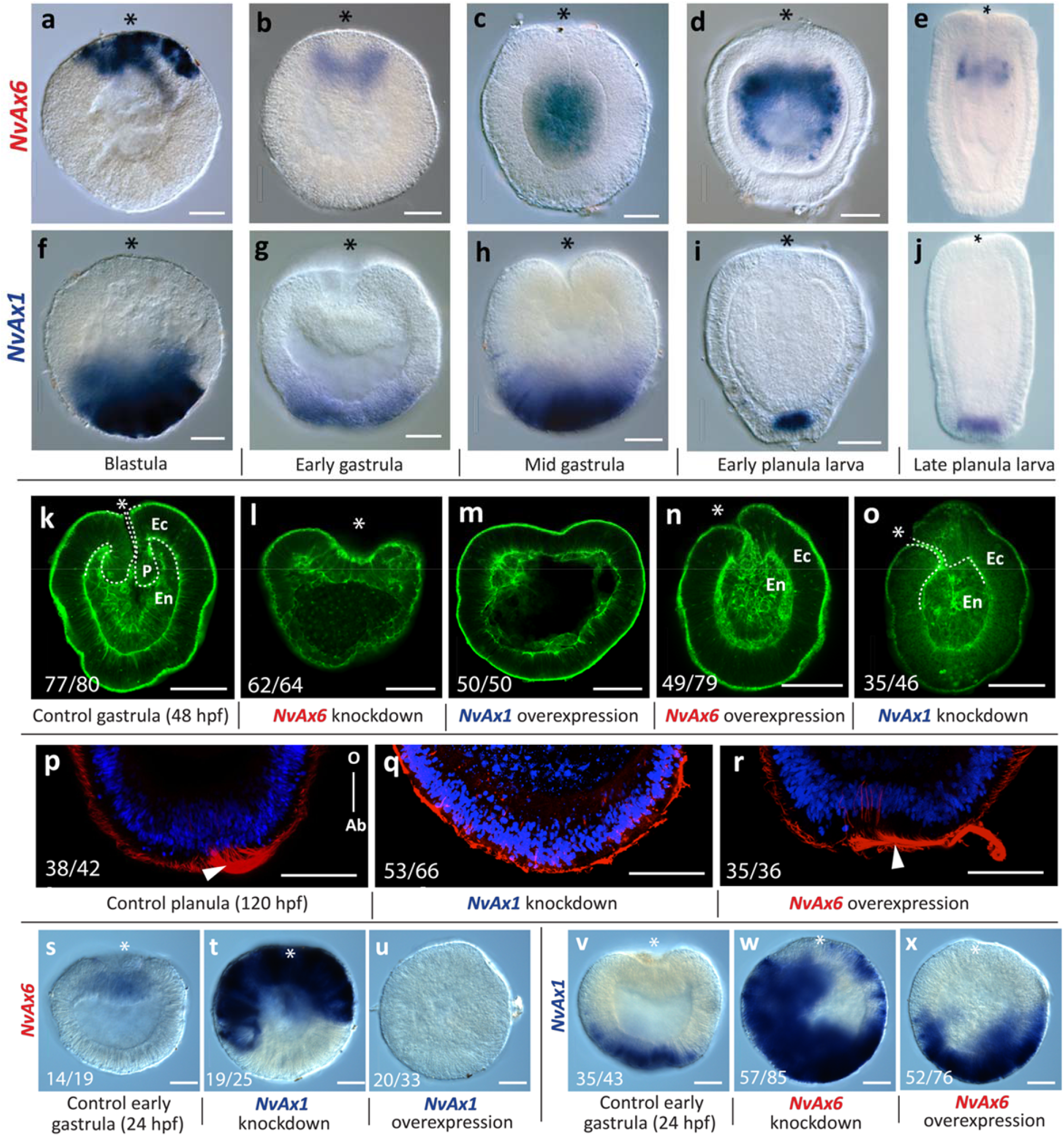
Perturbation of *Hox* expression disrupts oral patterning during gastrulation. Developmental time series of anterior, *NvAx6* **(a-e)**, and central/posterior, *NvAx1* **(f-j)**, *Hox* mRNA expression at blastula **(a,f)**, early gastrula **(b,g)**, mid gastrula **(c,h)**, early planula **(d,i)** and late planula stages **(e-j; images from Ryan et al. 2007**). **(k-o)** Gastrulation defects due to disruption of anterior (*NvAx6*) and central/posterior (*NvAx1) Hox* expression through microinjection of antisense translation blocking morpholinos (knockdown) or *in vitro* transcribed mRNA (overexpression). **(k)** Fluorescent phalloidin-labeled embryo during final stages of gastrulation (48hpf) with distinct ectodermal(Ec) and endo-mesodermal(En) tissue layers delineated by the early pharynx **(P)** (white dashed line delineates the pharynx from endomesoderm). **(l)** Knockdown of anterior *Hox (NvAx6*) blocks invagination of the presumptive endomesoderm. **(n)** Overexpression of central/posterior *Hox (NvAx1*) mRNA also blocks gastrulation and produces gastrula stage embryos with reduced axial morphology. **(m)** Overexpression of anterior *Hox (NvAx6*) mRNA and **(o)** knockdown of the central/posterior *Hox* gene (*NvAx1*) disrupts pharynx development and produces ectopic oral tissue at the blastopore (white arrowhead). **(p-r)** Apical tuft cilia labeled with and acetylated tubulin antibody (red) with the nuclei counter stained with DAPI (blue) in planula stage embryos treated with control morpholino **(p)**, central/posterior *Hox* morpholino **(q)**, and anterior *Hox* morpholino **(r)** (dextran control listed in Supplemental Fig. 3). **(s-u)** *In situ* hybridization of anterior *Hox (NvAx6*) at 24hpf in control **(s)**, central/posterior *Hox* knockdown **(t)**, and central/posterior *Hox* overexpression treatments **(u)**. **(v-x)** *In situ* hybridization of central/posterior *Hox (NvAx1*) at 24hpf in control **(v)**, anterior *Hox* knockdown **(w)**, and anterior *Hox* overexpression treatments **(x)**. Scale Bars are 50um.

### Oral and aboral *Hox* genes have opposing but not symmetrical roles in oral-aboral patterning

*NvAx6* and *NvAx1* are expressed in opposite territories throughout development and were manipulated by microinjection of uncleaved zygotes to test the functional role during development. Gene-specific antisense morpholino knockdown of the orally expressed anterior *Hox* gene, *NvAx6*, results in defective in gastrulation. These embryos initially form an inner endomesodermal plate (Fig. 3l), characterized by the highly reduced expression of the endomesodermal marker *NvSnailA* (Fig. 4a), but fail to continue gastrulation. Additionally, knockdown of *NvAx6* results in the loss of the pharyngeal marker *NvFoxA* and the oral marker *NvBrachyury* that is expressed at the ectodermal/pharyngeal boundary (Fig. 4a), and has been shown to regulate pharynx formation in anthozoans^47,48^. Control injections of dextran or control morpholinos does not impair expression of any genes analyzed herein (Supplemental Fig. 3). *NvAx6* knockdown shows minor expansion of aboral markers such as *NvFGF2A, NvSfrp1/5, NvDkk1/2/4*, and *NvSix3/6* (Fig. 4b), but does not appear to effect aboral development, as embryos treated with *NvAx6* anti-sense morpholino eventually form a swimming planula larva with an apical tuft at the aboral pole (Supplemental Fig. 4a-f).

**Figure 4.**
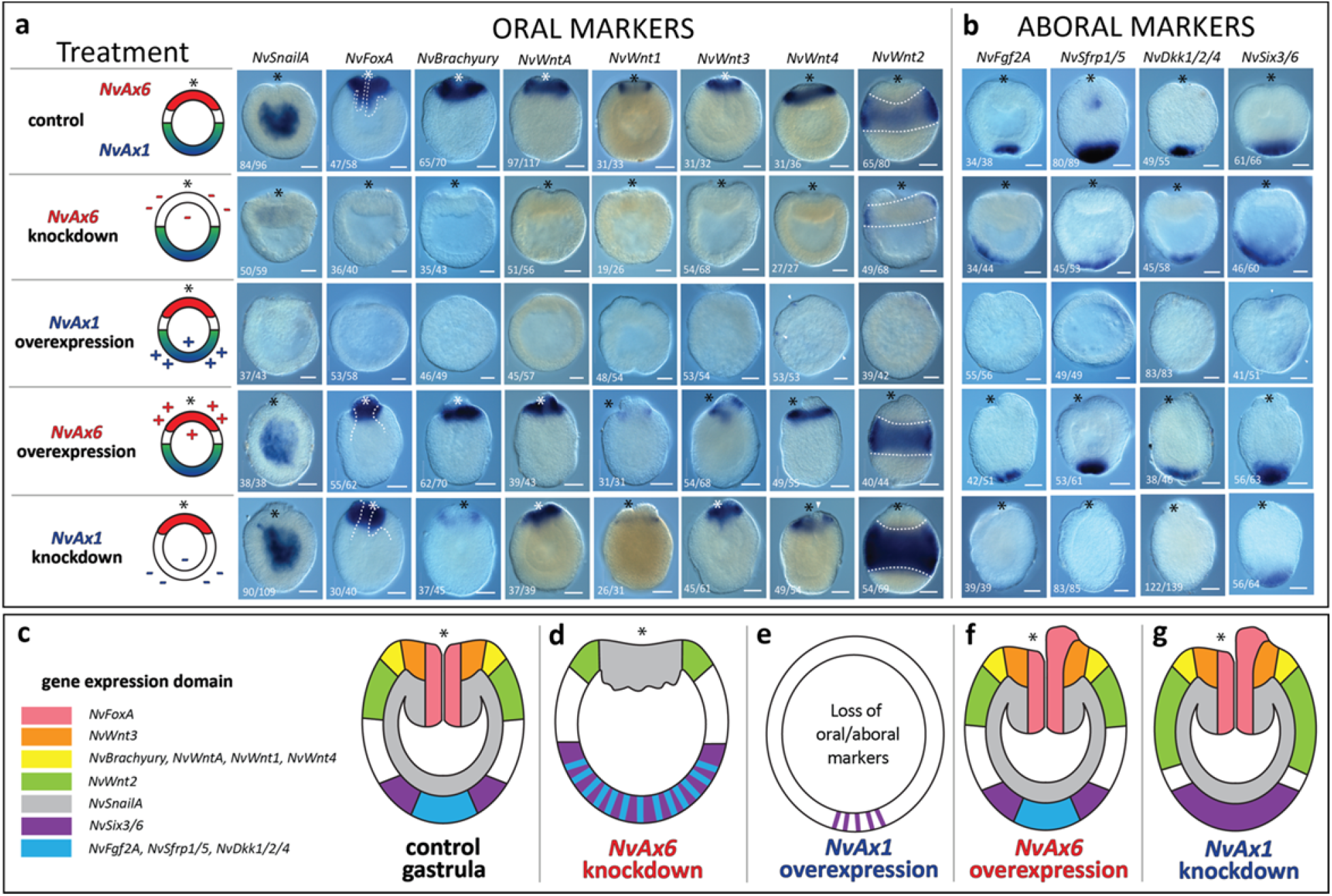
Anterior and central/posterior *Hox* genes have reciprocal phenotypes during oral but not aboral development. **(a)** Schematic representation of perturbation of either anterior *Hox (NvAx6*) or central/posterior *Hox* (Ax1) during early development. (KD=knockdown, ME=misexpression). **(a-b)** Late gastrula (48hpf) stage expression of molecular markers for oral **(a)** or aboral **(b)** territories. White dashed lines in *NvFoxA* column serve to highlight the larval pharynx and distinguish between the ectoderm and endoderm. White dashed lines in *NvWnt2* column outline the belt of *NvWnt2* expression. **(e-i)** Illustration summarizing territorial changes due to manipulation of Anterior (*NvAx6*) and central/posterior (*NvAx1) Hox* genes. Red and blue boxes represent respective oral and aboral territories. Scale Bars are 50um.

Overexpression of the aborally expressed *Hox* gene, *NvAx1*, mRNA results in a loss of gastrulation (Fig. 3m), with the absence of the endodermal plate or any morphological signs of invagination (Supplemental Fig. 4c), as well as the loss of *NvSnailA, NvFoxA* and *NvBrachyury* gene expression (Fig. 4a). Surprisingly, the apical tuft does not form (Supplemental Fig. 4f), and aboral markers are reduced or lost as a result of *NvAx1* overexpression (Fig. 4b). *NvSix3/6* appears both diminished and disorganized, expanding into multiple regions of the embryo (Fig. 4b).

Overexpression of the anterior Hox gene *NvAx6* or morpholino knockdown of *NvAx1*, each interfere with pharyngeal patterning. Both treatments produce external tissue with a pronounced asymmetry at the blastopore that fails to invaginate (Fig. 3n-o) and expresses the pharyngeal/oral markers *NvFoxA* and *NvBrachyury* (Fig. 4a). *NvSnailA* is expressed normally in the endomesoderm (or mesoderm^49^ ?) indicating that these defects are related to pharyngeal patterning and not endomesoderm specification (Fig. 4a). The existence of evaginated/ectopic pharyngeal tissue becomes even more pronounced in embryos injected with *NvAx6* mRNA that survive to later polyp stages (Supplemental Fig. 4b,h), but not in *NvAx1* knockdown treatments, which produced larvae without an apical tuft^50^ (Fig. 3q) and polyps with a greatly elongated body column and reduced oral morphology (Supplemental Fig. 4g,j). Aboral markers are lost in *NvAx1* knockdown treatments, suggesting that the *NvAx1* expression is necessary for aboral specification (Fig. 4b). However, aboral marker expression and apical tuft formation are not disturbed by *NvAx6* overexpression (Fig. 4b).

At early gastrula stages (24hpf), knockdown of the oral *Hox* gene, *NvAx6*, and the aboral *Hox* gene, *NvAx1*, caused an expansion of *NxAxl* and *NvAx6* expression respectively (Fig. 3t,w), pointing to a mutually antagonistic relationship. However, overexpression of the anterior Hox gene *NvAx6* exhibits only a slight expansion of *NvAx1* expression (Fig. 3x) but has no effect on apical tuft formation (Fig. 3r), while overexpression of aboral *Hox* gene, *NvAx1*, completely abolishes *NvAx6* expression at the oral pole (Fig. 3u) and inhibits oral development (Fig. 3u). This suggests that *NvAx6* and *NvAx1* maintain respective oral and aboral expression domains through mutual antagonism.

### Role of *NvAx6* and *NvAx1* in oral-aboral patterning during early development

Disruption of oral or aboral Hox genes expand opposing territories (Fig. 3t,w). Ectopic aboral *Hox (NvAx1*) expression restricts gastrulation and oral specification, resulting in the lack of endomesoderm formation throughout development (Fig. 4a). Similarly, knockdown of the oral *Hox* gene *NvAx6* produces a severe defect in gastrulation (Fig. 3l) and results in an expansion of *NvAx1* expression toward the oral pole (Fig. 3w). This suggests that stalled gastrulation as a result of *Hox NvAx6* knockdown is, in part, a result of the upregulation and expansion observed by *NvAx1* transcript towards the oral pole. To test this hypothesis, we co-injected both *NvAx6* (oral) and *NvAx1* (aboral) morpholinos into *N. vectensis* zygotes and assessed changes in late gastrulae (48hpf). Controls were injected with either *NvAx6* or *NvAx1* morpholinos with a standard control morpholino and produced phenotypes identical to those seen with normal single gene-specific morpholino injections (Fig. 5). Embryos treated with both *NvAx6* and *NvAx1* morpholinos undergo gastrulation to form an outer ectoderm (Ec) and an inner endomesoderm (En) that expresses the maker *NvSnailA* (Fig. 5), but fail to form a pharynx(P) or express the pharyngeal markers *NvFoxA and NvBrachyury* (Fig. 5). Aboral markers assessed in this study were lost in double injection (*NvAx6* + *NvAx1*) experiments (Fig. 5), including a complete loss of *NvSix3/6* (Fig. 5), which is repressed but not lost in *NvAx1* morpholino treatments. Co-injection of *NvAx6* and *NvAx1* morpholinos has no effect on oral marker expression, including the pharyngeal markers *NvFoxA* and *NvBrachyury* at earlier gastrula stages (24hpf) (Supplemental Fig. 5), suggesting that patterning cues downstream of the oral *Hox* gene, *NvAx6*, are required after the onset of gastrulation. However, the aboral marker *NvSix3/6* is lost at 24hpf in treated embryos (Extended Data Fig. 5), thus *NvAx1* expression has a patterning role at these earlier gastrula stages.

**Figure 5.**
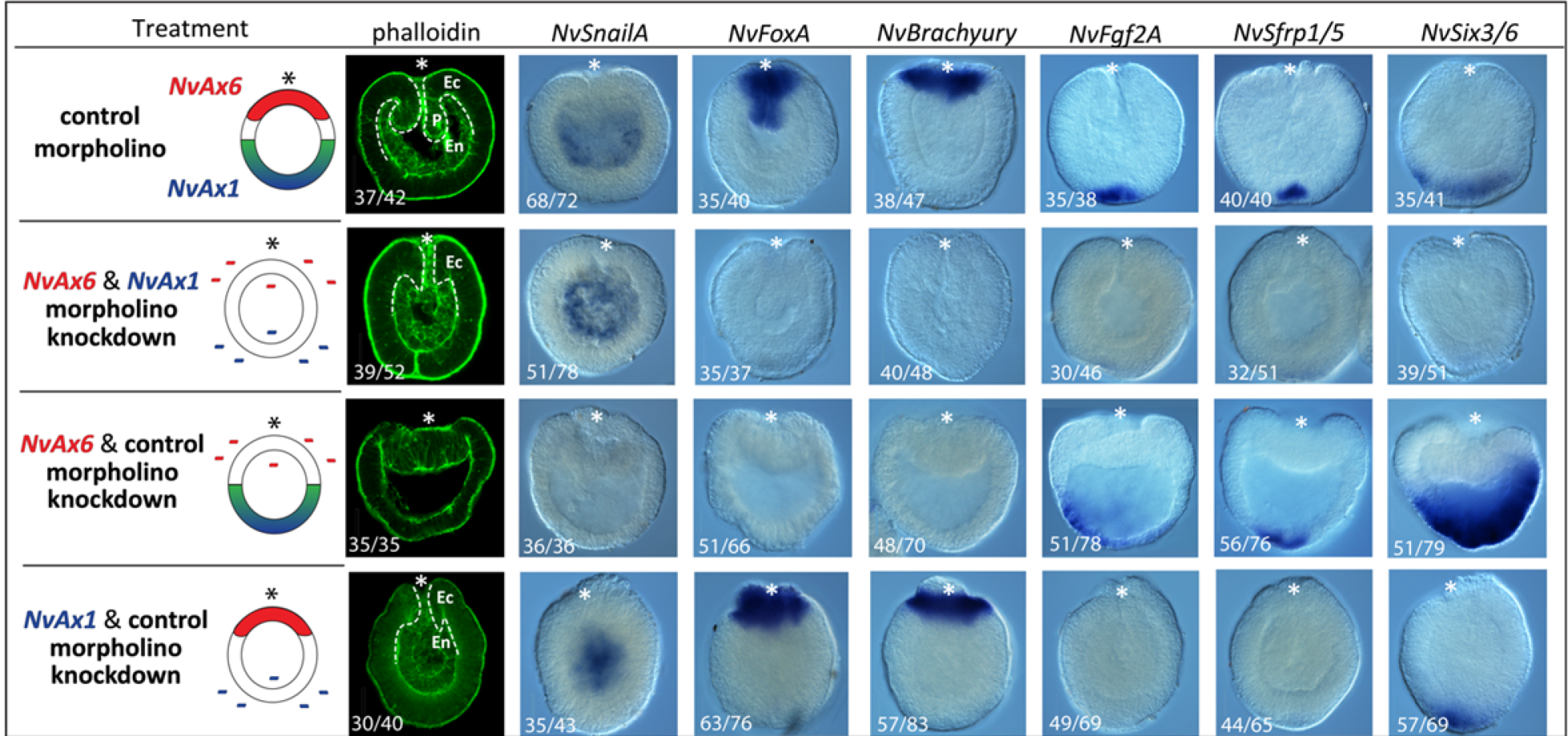
Loss of anterior and central/posterior *Hox* results in loss of oral-aboral patterning, but not endomesoderm formation. Gastrulation defects resulting from co-injection experiments as assessed by fluorescent phalloidin labeling (far left column) and *in situ* hybridization for oral (*NvSnailA, NvBrachyury* and *NvFoxA*) and aboral (*NvFgf2a, NvSfrp1/5* and *NvSix3/6*) markers. Ec = ectoderm, En= endomesodermal and P=pharynx. White dashed line serves to distinguish the point of transition between the ectoderm and the endoderm. Scale Bars are 50um.

### *NvAx6* and *NvAx1* regulate oral-aboral axis through interactions with Wnt signaling

The oral-aboral axis of cnidarians is thought to be patterned by restricted Wnt domains beginning at the onset of gastrulation and progressing through larval development^51–56^. *NvWntl, NvWnt3, NvWnt4*, and *NvWntA* are expressed around the blastopore/oral pole and disappear following both oral *Hox NvAx6* knockdown and aboral *Hox NvAx1* overexpression (Fig. 4a). *NvWnt2* is expressed in a band along the ectodermal midline of the embryo and serves as an important marker for defining oral and aboral territories. Knockdown of *Hox NvAx6* expression results in shift of *NvWnt2* expression towards the oral pole (Fig. 4a) while overexpression of *NvAx6* has no observable effect on *NvWnt2* expression (Fig. 4). *NvWnt2* expression is lost in aboral *Hox (NvAx1*) overexpression treatments, while knockdown *NvAx1* resulted in a robust upregulation and aboral expansion of the *NvWnt2* domain (Fig. 4). A more pronounced phenotype was produced in CRISPR cas9 mediated knockout of *NvAx1*, resulting in upregulation of *NvWnt2* throughout the entire ectoderm and endomesoderm, and of *NvWnt1* in the endomesoderm (Supplemental Fig. 6).

## Discussion

### Towards a molecular blueprint for oral-aboral patterning

Since the first cnidarian genome became available^57^ it has become clear that much of the developmental “toolkit” to build a bilaterian is present in the cnidarian lineage. Although there are aspects of cnidarian *Hox* clusters that have diverged from the ancestral condition in virtually all lineages (e.g., loss or duplication of *Hox2* homologs, Supplemental Fig. 1), it appears that the common ancestor of cnidarians and bilaterians utilized *Hox* genes to pattern their primary (oral-aboral) body axis. If a *Hox* cluster is representative of the ancestral condition in Cnidaria, gene duplications and loss may have played an instrumental role in divergent metagenic lifecyles within the phyla.

There exists a longstanding argument to the original of anterior-posterior patterning outside of bilaterians. These results indicate that anterior and central/posterior *Hox* genes specifying oral and aboral territories, respectively, before the onset of gastrulation. Currently, this study corroborates fate mapping results that show that the cnidarian animal pole corresponds to the oral pole^58,59^. Furthermore, these data are contrary to idea that swimming directionality indicates anterior-posterior positionality within cnidarian, arguments based on transient larval characters (the apical ciliary tuft)^60–62^. These findings would suggest that the oral-aboral axis is homologous to the bilaterian anterior-posterior axis with the adult mouth corresponding to anterior in both cases. Further characterization of germ layer segregation^49^ and cell-type diversity among cnidarians is necessary to understand territorial homology between cnidarians and bilaterians.

In this report, we’ve characterized for the first time in any cnidarian embryo the existence of early domains of reciprocal *Hox* gene expression in the starlet sea anemone, *Nematostella vectensis*, that begin at blastula stages and extends through gastrulation and that the interaction of these genes are functionally relevant for early axis formation and tissue specification along the oral-aboral axis. *NvHox* and related homeodomain containing genes may exhibit temporal but not spatial colinearity during early stages of development in *N. vectensis*. Anterior and central/posterior-like *Hox* genes maintain oral and aboral expression domains through mutual antagonism. The *Hox* gene *NvAx1* that is expressed at the tip of the aboral axis through embryogenesis and larval development is functionally dominant over the anterior *Hox* gene *NvAx6*. Although suggested to be primarily a vertebrate phenomenon^39–41^, elements of the hierarchal prevalence among *Hox* genes appears to be functioning in *Nematostella*. Further investigation into the mechanism of this relationship and the hierarchal nature of the other *NvHox* genes will help determine if the paradigm of posterior dominance is truly conserved between bilaterians and cnidarians, and will help resolve if *NvAx6* exhibits any form of anterior dominance.

Through these studies, we can begin to understand the molecular basis of oral-aboral patterning in *Nematostella* and develop a theoretical model of cnidarian *Hox* function (Fig. 6). Oral and aboral *Hox* genes of *Nematostella* are expressed in opposite domains during early development, and their spatial patterning is important for the establishment of oral and aboral domains (Fig 6a). *NvAx6* and *NvAx1* act in an antagonistic fashion to pattern both oral and aboral territories, which appears to be mediated through interactions with Wnt signalling (Fig. 6b). During early cleavage stages, the stabilization of β-catenin and components of both canonical and PCP Wnt signaling pathways establish the site of gastrulation/future oral pole in *Nematostella*^66,67^ (Fig. 6c; I). Recent findings have shown that *Nvβ-catenin* activity and Wnt signaling at the oral pole is necessary for the onset of aboral specification, a process that is partially mediated by the homeobox gene *NvSix3/6*^68^ (Fig. 6c; II). Our findings suggest that the maternal aboral marker *NvAx1* suppresses activation of *Wnt* signaling at the oral pole when overexpressed due to its inhibition of *Nvbra* (Fig. 6c; IV), in turn effecting both oral and aboral specification. Moreover, this explains the seemingly paradoxical results which suggest that endogenous *NvAx1* expression is a promoter of aboral development while ectopic *NvAx1* expression disrupts aboral specification, resulting from the restriction of Wnt-signaling at the oral pole. Further investigation into the functional relationship between aboral *Hox (NvAx1)* expression and Wnt signaling is required to validate this hypothesis. While this example occurs at a markedly later stage in embryonic development compared to the early stages of gastrulation assessed in this study, the formation of drastically elongated polyps in aboral *Hox (NvAx1*) knockdown (Supplemental Fig. 4j) points to a captivating similarity between the functions of vertebrate and anthozoan central/posterior (and posterior-like) *Hox* genes in axial elongation. Furthermore, results from the double knockdown of *NvAx6* and *NvAx1* provides support for the proposed role of *NvAx1* in restricting oral specification, possibly through the restriction of Wnt signaling, as parallel knockdown of *NvAx1* was sufficient to rescue the loss of gastrulation and oral specification phenotype produced in *NvAx6* morpholino treatments. From these results, we propose that *NvAx1* is an inhibitor of Wnt/β-catenin signaling (Fig. 6c; IV) similar to what has recently been described during vertebrate development^69^, and that the core regulatory relationship between central/posterior (posterior-like) *Hox* expression and Wnt signaling may be a highly conserved characteristic of posterior *Hox* function.

**Figure6:**
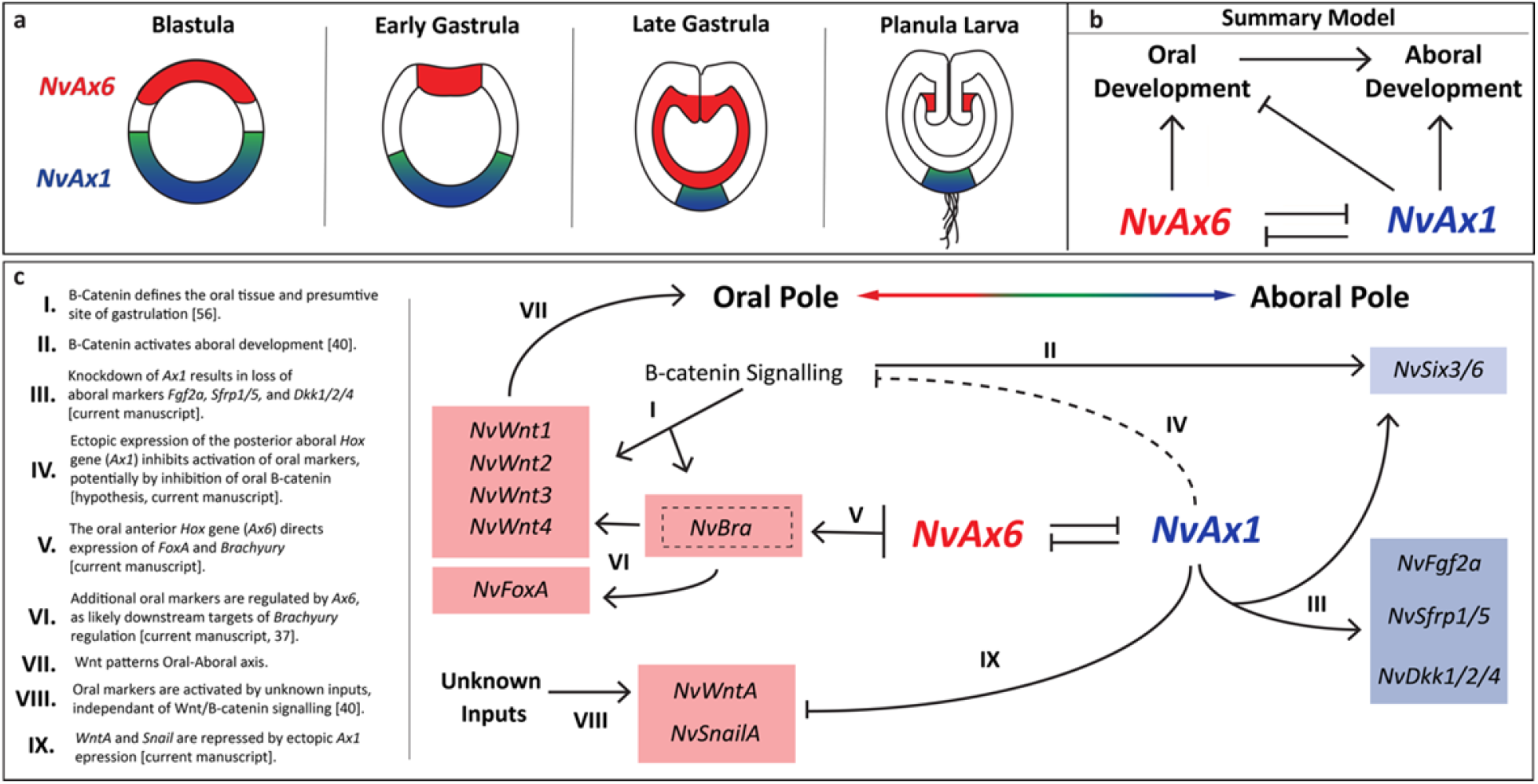
Model of *Hox* gene function in early development of an anthozoan. **(a)** Illustration depicting anterior *Hox (NvAx6*) (red) and central/posterior *Hox (NvAx1*) (blue) expression throughout early development of *N. vectensis*. **(b)** Working model of how anterior (*NvAx6*) and central/posterior (*NvAx1) Hox* genes function to specify the oral-aboral axis before and during gastrulation in an anthozoan cnidarian. **(c)** Illustration of known regulatory relationships between *Hox* genes and the oral (*NvWntA, NvWnt1, NvWnt2, NvWnt3, NvWnt4, NvFoxA, NvBrachyury*, and *NvSnailA*) and aboral (*NvSix3/6, NvFgf2A, NvSfrp1/5*, and *Dkk1/2/4)* markers assessed in this study. Important components of this regulatory paradigm needing description are marked with roman numerals (I-IX) with corresponding text at the periphery of the illustration. Regulatory relationships derived from previous studies are cited in the periphery text (I, II, VI, and VIII).

The phylogenetic origin of the *Hox* patterning system is a highly controversial matter among evolutionary developmental biologists. The cnidarian clade has the greatest diversity of *Hox* genes among the four basal animal lineages (sponge and ctenophore genomes do not contain *Hox* genes^33-35,37^). Genomic resources from a diverse group of cnidarians suggest rearrangements and fragmentations of genomic clusters are present among cnidarian species. Although many have argued that cnidarian *Hox* genes do not play a role in axial patterning^19,29,44^, this functional analysis of *Hox* genes in *Nematostella* suggest that some aspects of genomic organization and anterior-posterior patterning were present in the cnidarian ancestor. It will be interesting if the study of other cnidarian species reveal additional roles for *Hox* genes during early development, which will ultimately help resolve the shared and derived characters of cnidarian and bilaterian *Hox* genes. Together these data suggest that cnidarians retain components of a simple ancestral *Hox* system that was originally deployed during early developmental stages and functioned to organize the primary axis in the cnidarian ancestor.

## Materials and Methods

### Animal Care

Adult *Nematostella vectensis* were raised in 1/3x Seawater at 16 °C in constant dark. Animals were fed artemia once a week and fed oyster forty-eight hours before spawning, which was engendered by an eight hour light box cycle at 16 °C. Distinct groups of animals were spawned once every 3-4 weeks. Fertilized embryos were collected and placed in a 4% cysteine wash (4%cysteine in 1/3 filtered seawater, pH7.4) for fifteen minutes to remove the outer jelly layer. Eggs were then washed 3x with 1/3 filtered seawater before conducting experiments. Treated embryos were raised to collection at 16°C. Embryos used were from randomized individuals from different genetic individuals to eliminate genetic variability. Mixed genetic pools of individual animals created a blind test of the effectiveness of each treatment regardless of genetic ancestry.

### Microinjection: knockdown, overexpression and knockout

We disrupted *NvAx6* and *NvAx1 Hox* gene expression using gene-specific antisense translation blocking morpholinos or CRISPR/Cas9 mediated genome editing using multiple guide RNAs against each gene^47,70,71^. Genes were over expressed by injecting *in-vitro* transcribed mRNA^72^. Morphological and molecular analysis of phenotypes was conducted during late gastrula (48hpf), planula larvae (96hpf) stages of development and 1-2 week old polyps. Control injections using a standard control morpholino and dextran lineage tracer (or Cas9 protein without guide RNAs in knockout injections) produced normal gastrula stage embryos (Fig. 3k) possessing an outer ectoderm(Ec), inner endomesoderm(En), and pharynx(P) (dashed white line) (Supplemental Fig. 3). Knockdown and knockout treatments produced similar molecular phenotypes (Supplemental Fig. 6), thus validating the efficacy of the *NvAx6* and *NvAx1* morpholinos. Morpholino and mRNA injections were conducted following previously described methods^72^. These methods were further adapted for CRISPR/Cas9 injections (described below). Treated embryos were raised to collection at 16°C.

Translation blocking morpholinos against *NvAx6* and *NvAx1* and a standard control morpholino have been developed through Gene Tools, LLC. Philomath OR, 97370. Morpholino sequences are listed in (Standard Control 5’-CCTCTTACCTCAGTTACAATTTATA -3’; *NvAntHox6* 5’-ACCGCCGCTCATGCCCAAATGTGTC -3’; *NvAntHox1* 5’-TTGACTGCATGATGTGCGCTCTAGT -3’). Expression constructs for *NvAx6* and *NvAx1* were generated using the gateway cloning system (SPE3-Ax1-Rvenus, SPE3-Ax6-RmCherry and SPE3-Ax6a-RmCherry). Cloned sequences are listed in (Supplementary Data Table 1). Morpholinos and mRNA were first injected in a dilution series ranging in concentration from 0.1mM and 0.9mM and appropriate concentrations (0.3mM for *NvAx6* MO, 0.5mM *NvAx6* mRNA, 0.9mM for *NvAx1* and 0.3mM *NvAx1* mRNA) were selected to achieve an optimal relationship between toxicity and phenotype penetrance.

CRISPR/Cas9 genome editing in Nematostella was performed s previously decribed^47,73^. Briefly, target sequences 18-20bp in length fitting the guide-RNA (gRNA) target profile of 5’-G(G-A)-N(16-18)-NGG-3’ were identified within the coding sequence of the *NvAx6* and *NvAx1* locus (Supplemental Fig. 7a & 8a) using a web-based program called ZiFiT (http://zifit.partners.org). Six target sequences were selected for each locus bases on their location (choosing sites scattered throughout the coding sequence) and their gRNA efficiency score, which was calculated using the web-based CRISPR Efficiency Predictor provided by the DRSC at Harvard Medical School (http://www.flvrnai.org/evaluateCrispr/). Sequences and efficiency scores are listed in (Supplementary Data Table 1). Additionally, potential target sequences were blasted against the publically available *Nematostella* genome (http://genome.jgi.doe.gov) to insure that there were minimal off-target sites related to the sequence, limiting partial off-target sites to having no more than 15 base pairs in common with the 18-20 base pair target sequence. Oligonucleotides were generated integrating the target sequence (5’-G(G-A)-N(16-18)-NGG-3’) into a CRISPR RNA (crRNA) sequence containing a T7 (5’-AATTAATACGACTCACTATA -3’) or Sp6 (5’AATATTTAGGTGACACTATA 3’) promoter. Full-length gRNA was generated following a previously published protocol^74,75^, starting with PCR assembly with a trans-activating crRNA (tracrRNA) oligonucleotide (Supplemental Table 2), followed by *in vitro* transcription (NEB Highscribe™ T7 and Sp6 RNA synthesis kits; Cat# E2040S and E2070S), and RNA purification (Zymogen RNA Clean and Concentrator™-25; Cat# R1017). gRNAs at a final concentration of 400ng/ul (consisting of equal concentrations of each gRNA) were injected with Lyophilized bacterial type II Cas9 protein (PNA Bio, Thousand Oaks, CA) (1ug/ul) reconstituted in 50% glycerol and 0.1 mM DTT to generate frame shift mutations and deletions within the target locus. Genomic DNA was isolated from 8 individual 18hpf embryos per treatment group for PCR analysis of the targeted region following a previously published methods^74,75^. (Supplemental Fig. 7b & 8b). Additionally, *in situ* hybridization was performed on treated blastula stage embryos to ensure that gene expression was lost (Supplemental Fig. 7c & 8c). Treated embryos were collected at 48hpf for in situ hybridization studies to assess changes in marker gene expression (Supplemental Fig. 4).

### Quantitative PCR

Quantitative PCR was performed using the Roche LightCycler^®^ 480 Instrument II and LightCycler 480 SYBR Green I Master mix (Cat# 04707516001, Roche, Inc.). qPCR samples were standardized with *NvGADPH* and *NvRiboPro* and primers for other genes were designed using MacVector (www.macvector.com) to amplify 75-150 base-pair fragments of the desired gene. These primers were then back-blasted against the *Nematostella* genome to make sure they only will amplify a single region from the genome. We checked each primer efficiency with a dilution curve (10^-1^-10^-5^) to make sure their range was within the negligible value of 1.9-2.0. Individual stages were collected from pooled samples of embryos consisting of roughly 100 embryos. These stages represent a single biological sample, and were confirmed through technical replication. Total RNA from each sample was stored in TRIzol (15596-026) at -80**°**C until processed. RNA processing and cDNA synthesis has been previously described^25^. Relative fold change values were calculated in Microsoft Excel and were standardized against our reference genes based on published formulas^75^.

### In situ hybridization

All *in situ* hybridizations were based off of the previous protocol for *Nematostella vectensis^76^*. Fixations were done in 1% gelatin coated dishes to prevent tissue from sticking to the plastic (sticking to plastic causes tissue damage and non-specific staining). Embryos were fixed in ice cold 4% paraformaldehyde with 0.2% glutaraldehyde in 1/3x seawater for two minutes, followed by 4% paraformaldehyde in 1/3x seawater for one hour at 4°C. Probe sequences (Supplementary Data Table 2) ranging from 550-1200 bps were cloned from cDNA using the pGEM^®^-T vector system and DIG-labeled RNA probes were generated following an established published protocol^76^. If available, cloned marker gene sequences were selected from a probe stock originally created for a previous study^46^. Probes were hybridized at 64°C for two days and developed with the enzymatic reaction of NBT/BCIP as substrate for the alkaline phosphatase-conjugated anti-DIG antibody (Cat.# 11093274910, Roche, Inc.). Wild type samples were developed for an equal amount of time and if no expression was visible, a subset of samples remained in developing solution for at least 1 day to determine if lower levels of expression was present. When developing functional in situs, development time was based on the signal seen in control samples. All experiments exhibited greater than 75% penetrance of phenotypes. Due to variability during injection, embryo size and development time of different genes, greater than 75% in replicates was considered the dominant phenotype created by genetic manipulations. Furthermore, to obtain material to test all the genes, samples were collected from multiple days of injection and pooled for in situ hybridization. Replicate attempts of in situ hybridization were attempted at least twice on pooled samples to obtain sample sizes generally greater than thirty embryos per replicate.

### Immunostaining

Methods for Immunostaining were adapted from previously published protocols^51^. Embryos collected for immunostaining were fixed in ice cold 4% paraformaldehyde with 0.2% glutaraldehyde in 1/3x seawater for two minutes, followed by 4% paraformaldehyde in 1/3x seawater for one hour at 4°C. Samples were then washed five times (5 min. each) in PBT (1% Triton-x and 1% BSA in PSS) and stored in PBS at 4°C for up to one month. Samples were then washed three times (30 min. each) in PBT followed by a one hour block in blocking solution (5% normal goat serum (NGS) in PBT) at room temperature. Samples were then treated with the primary antibody (monoclonal anti-α-Tubulin; SigmaT9026) diluted 1:500 in blocking solution and incubated overnight (12 hours) at 4°C. Primary antibody was removed and samples were rinsed five times (10 min. each) in PBT before adding the secondary antibody (goat-anti-mouse-568; A-11004) diluted 1:250 in blocking solution and incubated over night at 4°C. Tissues were then washed at least three times (15 min. each) in PBT before adding counterstain. DAPI was used (0.1ug/ul in PBS) was used to label cell nuclei. Fluorescent phalloidin (Invitrogen A12379) was used at a concentration of 1:200 in PBT to stain filamentous actin, incubating for no more than three hours at room temperature. Samples were washed at least three times (15 min. each) in PBS before clearing. Samples were cleared in 80% glycerol or using Murray’s Clear (1:1 benzyl benzoate and benzyl alcohol).

### Microscopy and imaging

Results from in situ hybridization studies were imaged using a Zeiss Axio Imager Z1 with a Zeiss HRc color digital camera run by Zeiss Zen 2012 software. Fluorescently labeled embryos were imaged using a Zeiss LSM-710 confocal microscope. Images of polyp stage results and whole wells of 24 well plates (as pictured in supplemental figures 4 and 5) were imaged using A Zeiss SteREO Discovery.V8 stereoscopic microscope with a Cannon EOS 5D MarkII full frame DSLR camera. Images were processed and scale bars were added using the Fiji distribution of ImageJ (http://fiji.sc).

### Data availability

Full length transcripts used for probe production, mRNA misexpression and designing morpholinos are available in the supplementary information.

